# Bloom syndrome helicase is required for efficient HIV-1 reverse transcription in macrophages

**DOI:** 10.64898/2026.04.26.720894

**Authors:** Andrew A. Leal, Samantha Tafrate, Xianbao He, Daniel L. Bryant, Melissa B. Herring, Maria Andrade, Joseph B. McWhirter, Andrés A. Quiñones-Molina, Suryaram Gummuluru, Manish Sagar, Rachel L. Flynn, Ignaty Leshchiner, Andrew J. Henderson

**Author notes:** Corresponding author: Andrew J. Henderson, Boston University Chobanian & Avedisian School of Medicine Department of Medicine and Virology, Immunology, Microbiology Section of Infectious Diseases, 650 Albany St, 6^th^ Boston, MA, 02118.

## Abstract

The induction of DNA damage by HIV-1 prior to integration suggests a function for DNA damage responses (DDR) during early infection, however what this role is remains incompletely understood. Initial experiments, using specific inhibitors for DDR pathways demonstrate that both ATM and ATR are necessary for efficient HIV-1 infection of macrophages with ATM acting at the late reverse transcription step. To identify DDR factors associated with ATM/ATR pathways that influence HIV-1 infection, a CRISPR knockout screen using a DDR-focused sgRNA library was performed. Approximately 30 DDR genes that impacted HIV-1 infection were identified with 13 factors that facilitated HIV-1 infection and 17 DDR factors that restrict HIV-1 infection. Several hits were factors associated with the Fanconi anemia pathway, such as BTR complex proteins, including the RecQ helicase Bloom syndrome helicase. BLM was specifically demonstrated to enhance HIV-1 infection and replication with knockdown of BLM expression diminishing integration and the establishment of intact HIV-1 proviruses in macrophages by 50%. BLM is associated with HIV-1 late reverse transcription intermediates, the step that proceeds HIV-1 integration. These findings identify BLM as a DDR host factor that promotes early HIV-1 infection by facilitating completion of reverse transcription and subsequent integration.

**Significance Statement:** HIV-1 infection elicits cellular DNA damage responses although the role of DNA damage in HIV-1 infection has not been fully characterized. Our study identifies specific DNA Damage factors that facilitate or restrict HIV infection. In particular, we show the RecQ helicase Bloom syndrome helicase (BLM) is a mediator of HIV-1 reverse transcription and integration in macrophages. This work highlights a functional interface between DNA damage repair pathways and HIV-1 integration suggesting that targeting select host DNA damage response factors can limit HIV-1 infection and persistence.

## Introduction

HIV-1 eradication has proven challenging largely due to the establishment of latent viral reservoirs. Generation of both latent reservoirs and actively transcribing proviruses occur after conversion of the viral RNA genome to double stranded DNA followed by insertion into the host cell genome (1–3). Despite suppressive antiretroviral therapies, these persistent integrated HIV-1 proviruses persist in cells presenting a major barrier to cure.

Orchestration of HIV-1 reverse transcription includes multiple template switches and generation of noncanonical RNA/DNA and single stranded/double stranded DNA structures (4). This process is mediated by the reverse transcription complex (RTC) that includes viral reverse transcriptase (RT), nucleocapsid (NC), and integrase (IN). Several putative and suspected cellular host factors are associated with the RTC that influence reverse transcription. Factors including the RNA helicase DHX9 and DNA topoisomerase TOP1 independently enhance RT activity (5–8). In contrast, increased levels of the RNA helicase MOV10 restricts the first and second strand transfers of reverse transcription (9, 10).

HIV-1 integration requires recruitment of several host proteins. The preintegration complex (PIC) associates with the completed reverse transcribed HIV-1 cDNA. The PIC is comprised of viral proteins IN, MA, NC and viral accessory protein Vpr, as well as host proteins that together mediate effective integration (11). Many of the PIC-associated host proteins are factors important for cellular DNA damage responses (DDRs) including LEDGF/p75, which is crucial for tethering the PIC via IN binding, to the chromatin sites of integration (12, 13). In addition, the RNA helicase AQR was shown to facilitate HIV-1 integration into R-loop dense genomic regions (14). The mechanism of integration induces DNA damage that host DDR factors are responsible for repairing (15). Furthermore, HIV-1 infection is associated with an induction of DNA damage and cell cycle arrest mediated by Vpr (16, 17). These observations suggest that cellular DDRs are critical for HIV-1 infection, and that HIV-1 co-opts these processes to assure efficient RT and provirus integration.

The DDR consists of networks of proteins that recognize and respond to exogenously or endogenously induced DNA lesions. Central to these networks are transducing proteins that propagate signal cascades to recruit and activate downstream effector proteins (18). Important transducers include three kinases from the phosphoinositide-3-kinase-related protein kinase family (PIKK), ataxia-telangiectasia mutated (ATM), ataxia-and Rad3-related (ATR), and DNA-dependent protein kinase (DNA-PK) (19). ATM and DNA-PK primarily respond to double-strand DNA (dsDNA) breaks, while ATR responds to single-strand DNA (ssDNA) breaks and replication stress (20). Another key transducer is Poly(ADP-ribose) polymerase 1 (PARP-1), which senses ssDNA breaks and influences specific types of DNA repair (21). Downstream DDR factors mediate multiple cellular responses including cell cycle arrest, chromatin remodeling, DNA replication fork stabilization, DNA repair, or apoptosis and cellular senescence. It remains unclear how host DDR proteins interact with early HIV infection and the extent they negatively or positively influence successful infection.

We employed functional genomic, epigenomic, pharmacologic and biochemical approaches to explore the intersection of HIV-1 and the DDR. We establish that ATM, ATR and PARP-1 inhibition in primary monocyte-derived macrophages (MDMs) negatively affect infection at different steps of the HIV-1 replication cycle. Using a DDR-specific CRISPR/Cas9 screen, we identify DDR factors that influence HIV-1 infection. BLM, a critically important and multifaceted RecQ helicase, was identified to facilitate HIV-1 infection. BLM knockdown in MDMs reduced total integrated HIV-1 and increased defective proviral integration due to diminished late reverse transcription. Importantly, we show BLM binds HIV-1 DNA during reverse transcription, before integration. Our study provides insights into host DDR/HIV-1 interactions and reveals a role for BLM helicase in mediating efficient HIV-1 infection.

## Results

### ATM and ATR inhibition reduces HIV-1 infection at different steps of the viral replication cycle in macrophages

Previous work addressing the roles of DNA-damage response pathways in HIV-1 infection have provided an incomplete and conflicting picture (22–25). PIKK transducers play pivotal roles in HIV-1 infection, but limited work on DDR pathways has been done in terminally differentiated or non-cycling myeloid cells, which are potential tissue-resident reservoirs for HIV-1 (26, 27). Studies exploring the roles of DDR protein PARP-1 in HIV-1 infection are discordant. Some data indicate that PARP-1 facilitates integration and transcription of HIV-1 while other groups have shown no effects on efficient integration and repression of viral transcription (28–31). We sought to determine if inhibiting key DNA-damage response pathways significantly altered HIV-1 infection in primary monocyte-derived-macrophages (MDMs). Cells were pretreated with drugs that potently inhibit ATM (KU-55933) (32) and ATR (VE-821) (33) kinases, as well as PARP-1 (olaparib) (34). Infections were done using VSV-G pseudotyped replication competent HIV-1 NL4-3 BaL *env* to facilitate efficient virus entry in the first round of virus infection. Three days post-infection (p.i.), cell supernatants were analyzed for p24^gag^ content by ELISA (**Fig. 1A**). Inhibiting any of these pathways led to significant reduction in HIV-1 p24^gag^ secretion in cell supernatants. In particular, ATM and ATR inhibitors decreased HIV-1 replication by greater than 90%, whereas PARP-1 had a more modest impact, decreasing HIV-1 replication by 60%. These decreases in HIV-1 replication were not due to cell toxicity of compounds with cell viability following treatment with all inhibitors being approximately 90% for all (**Fig.S1**)., To identify the steps of HIV-1 replication the inhibitors were disrupting, we examined HIV-1 integration by a nested Alu-element proximal PCR (Alu-PCR) two days p.i. (35). There were significant reductions of HIV-1 proviral integration in ATM and ATR inhibited cells, with ATM inhibition resulting in greater than a 75% decrease in proviral copy numbers (**Fig. 1B**). We also measured 2-LTR circles by qPCR to determine if there was a disruption in episomal HIV-1 DNA formation and observed reductions of greater than 80% with ATM inhibition and 50% with ATR and PARP (**Fig. 1C**). Using MDMs from five different people there was a wide range of 2-LTR circles measured in cells treated with the ATR inhibitor, possibly reflecting donor variation. To examine if a specific step of reverse transcription was being inhibited, RT early, intermediate and late products were measured by qPCR. ATM inhibition had the largest impact on late RT products reducing detection of these products by >50%, whereas the ATR and PARP inhibitors had a minimal effect on RT (**Fig. 1D-F**). These results implicate a role for DDRs, especially ATM, in facilitating early HIV-1 infection are consistent with some previous studies (22, 36–38).

**Figure 1.**
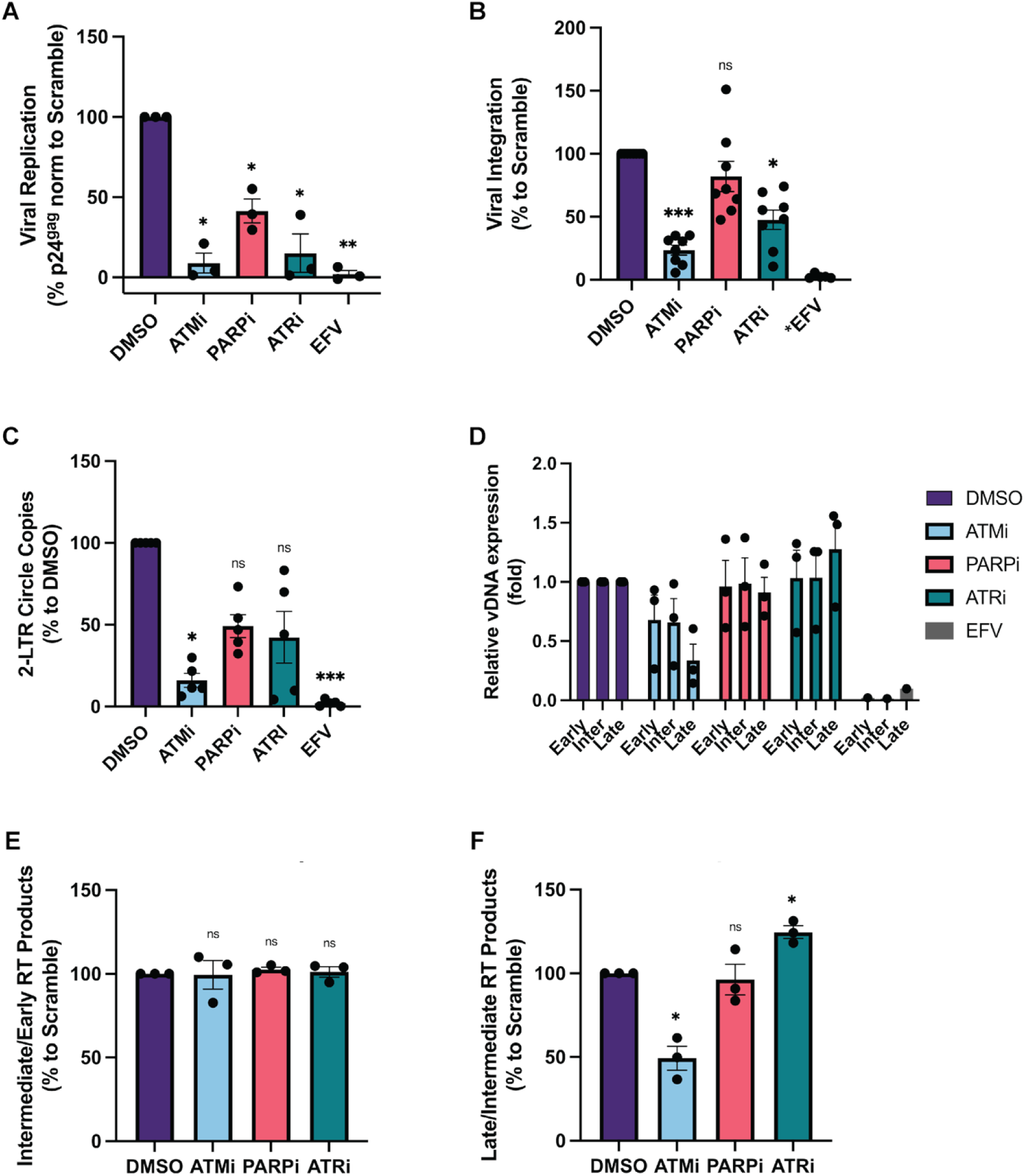
ATM/ATR inhibition reduces HIV infection at multiple steps in MDMs. MDMs were infected with NL4-3 BaL after pre-treatment with DMSO (10 μM), KU-55933 (ATM inhibitor, 10 μM), olaparib (PARP1 inhibitor, 10 μM), or VE-821 (ATR inhibitor, 10 μM) for 18 h. Controls included MDMs pre-treated with efavirenz (EFV, 1 μM) 1h before infection. Cells were post-treated and maintained on 5μM of DDRi drugs. **(A)** p24^gag^ was measured from cell supernatants with ELISA 3 dpi. **(B)** Total cell DNA was extracted 3 dpi and assed for proviral integration by Alu-PCR and **(C)** 2-LTR circle formation by qPCR. **(D)** DNA was extracted 2 dpi and assayed for RT products by qPCR. Early, intermediate (Inter), and late RT products were calculated compared to normalized untreated control infections (DMSO). **(E)** Percentage of intermediate RT products and **(F)** late RT products were determined by comparing ratios to untreated (DMSO) controls. Data are displayed as the means ± SEM with each point representing a different MDM donor. Statistical significance assessed by 1-way ANOVA with Dunnet’s multiple comparisons (to DMSO control) ns: not significant, *: p<0.05,**: p<0.01, ***: p < 0.001.

### DDR genes that influence HIV-1 infection identified using a CRISPR screen

Having established that inhibition of general DDR pathways negatively affects HIV-1 replication in macrophages, we aimed to identify specific host factors downstream of these kinases that influence HIV-1 infection. To identify DDR factors, we performed a CRISPR/Cas9 knockout screen by transducing immortalized human microglial cells (CHME3 cells) with a DDR-specific library of sgRNAs targeting 365 DDR genes (39). Guide coverage was maintained at 1000X throughout the experiment. Cells were infected with a single-round GFP-reporter HIV-1 for 2 days and sorted by flow cytometry for GFP-positive and negative populations. Following expansion for 5 days, these populations were analyzed by Next Generation Sequencing (NGS) to identify putative candidate genes. We measured the degree that guides were depleted or enriched relative to non-essential targeted gene controls and against GFP-negative or uninfected experimental conditions, by calculating the Model-based Analysis of Genome-wide CRISPR-Cas9 Knockout (MAGeCK) Robust Rank Aggregation (RRA) score for each gene (40) (**Fig. 2A**). A total of 137 genes that potentially altered HIV-1 infection were significantly enriched or depleted compared to controls (**Fig. S2, Fig. 2**). The two datasets corresponding to GFP-positive vs GFP-negative and GFP-positive vs no infection were compared and 32 genes overlapped between the two analyses **(Supplemental data sets 1 & 2, Fig S3, S4, Fig. 2B-D**). Seventeen gene knockouts (KOs) were enriched in GFP+ cells, suggesting they are antiviral or restrictive host factors to HIV-1 **(Fig. S4, Fig. 2C**). Thirteen different gene KOs were robustly depleted in GFP-positive cells, and thus expression of these genes were associated with decreased HIV-1 infection, suggesting pathways required for efficient infection **(Fig. S3, Fig. 2D**). Two genes, TOP1 and CDC25B, were discordant between analyses, being enriched in one dataset and depleted in the other. Multiple members of the Fanconi anemia pathway have been implicated as being involved in efficient HIV-1 infection and several of our hits, FANCI, FANCM, FANCB, FANCC and FANCG, were consistent with these previous studies (15, 41) (**Fig. 2D, Fig. S2-4**). Network analysis of the overlapping, significant hits from the screen using the STRING-db protein:protein interaction network revealed potential DDR complexes facilitating infection. The majority of hits identified in the depleted gene set are DNA repair factors. RMI1, RMI2, and BLM consist of three of the four BTR complex proteins. Other repair factors include FANCI, UIMC1 (RAP80), DDX11 and ETAA1 (**Fig. 2E**). Together these data from our CRISPR screen identified several putative DDR host factors that either facilitate or inhibit HIV infection.

**Figure 2.**
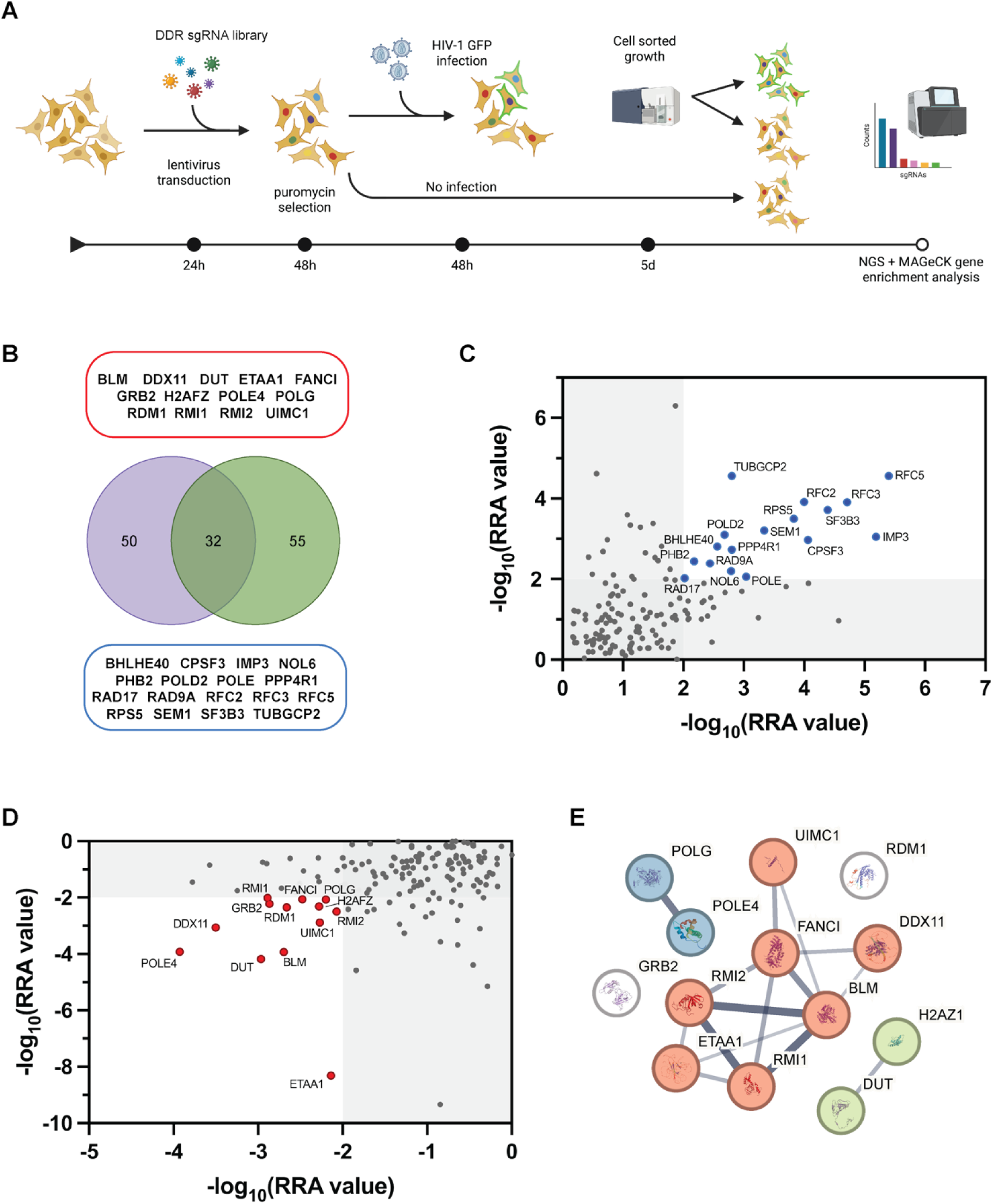
CRISPR screen identifies DDR genes influencing HIV-1 infection **(A)** DDR-specific CRISPR screen workflow. MAGeCK scores were calculated for genes sequenced in the GFP+ group compared to GFP- and compared to HIV-1 negative groups. **(B)** 32 genes overlapped when both analyses compared against each other. **(C)** A scatterplot comparing MAGeCK scores (-log(RRA value) of each analyses showing restrictive host factors to HIV-1 infection and **(D)** facilitative host factors for HIV-1 infection. **(E)** STRING-db protein:protein interactions using Markov Clustering (MCL) for facilitative genes

### BLM knockdown inhibits HIV-1 infection in macrophages

The RecQ helicase BLM was identified in our CRISPR screen as a facilitator for HIV-1 infection. BLM displays a diverse set of functions that are critical to genome stability including dissolution of Holliday junctions, G-quadruplexes, and D-loops as well as assisting in telomere DNA replication (42–44). BLM regulates double-strand break DNA repair and stabilizes replication forks by facilitating replication restart and suppressing new origin firing (45, 46). BLM complexes with RMI1, RMI2 and TOP3A forming the BTR complex and promotes DNA dissolution without crossover events (44). This complex is recruited by FANCM, through RMI binding, during Fanconi anemia DNA repair pathway activation (47). To explore if these components influence HIV-1 infection in macrophages, we knocked down proteins of the BTR complex, BLM, RMI1, and FANCM in MDMs using pooled siRNAs 6 days post-differentiation. Cells were then infected with HIV-1 NL4-3 BaL pseudotyped with VSV-G. Successful knockdowns were validated by diminished RNA expression of each of the targeted factors (**Fig. 3A**, **Fig. S5**). Initially, we assessed the impact of the BTR complex on HIV-1 replication by monitoring p24^gag^ in the supernatant with ELISA (**Fig. 3B**). Knockdown of BLM or FANCM expression resulted in significant ∼50% reduction in p24^gag^ secretion in MDMs compared to scrambled siRNA controls, whereas RMI1 knockdown had minimal impact on HIV-1 replication (**Fig. 3B**).

**Figure 3.**
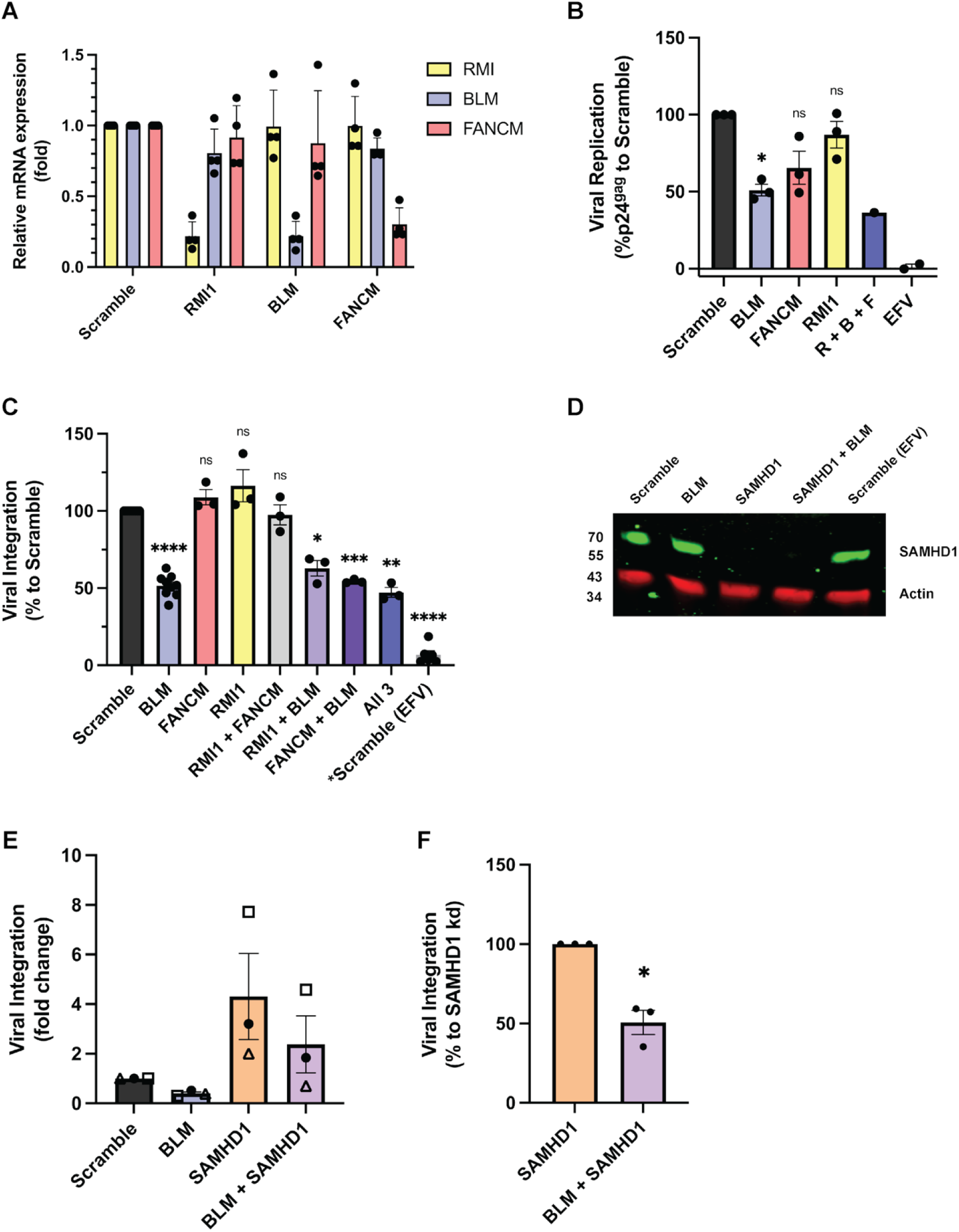
BLM knockdown inhibits HIV-1 infection in macrophages **(A)** mRNA expression for each targeted gene indicated was measured by qPCR following a double-treatment of 100 nM siRNAs in MDMs. **(B)** MDMs from 3 different donors were infected with NL4-3 BaL after pre-treament with siRNAs and 3 dpi, supernatants were measured for p24^gag^ by ELISA. **(C)** Proviral integration, 2 dpi measured by Alu-PCR. **(D)** siRNA knockdown of SAMHD1 was validated by western blot. **(E)** Proviral integration measured by Alu-PCR. **(F)** Paired donor experiments were normalized to SAMHD1 knockdown and expressed as a percentage of integration. Data are displayed as the means ± SEM with each point representing a different MDM donor. Statistical significance assessed by 1-way ANOVA with Dunnet’s multiple comparisons (to scramble siRNA control) or paired two-tailed t-test (to SAMHD1 siRNA). ns: not significant, *: p<0.05, **: p<0.01, ***: p < 0.001, ****: p < 0.0001.

To gain insights into the step of HIV-1 replication BLM and FANCM are acting at, we assessed HIV-1 integration by Alu-PCR in MDMs upon knockdown of BLM, FANCM or RMI1 (**Fig. 3C**). A significant ∼50% reduction of proviral integration was observed upon knockdown of BLM expression compared to scrambled controls. In contrast, no such reduction was observed in RMI1 or FANCM knockdowns, suggesting a different mechanism of action for the ability of FANCM to influence HIV-1 replication than that of BLM. We did not observe any additional impact on HIV-1 integration with knocking down multiple BTR factors and the 2-fold reduction of integration was only observed in cells that BLM was reduced. These data suggest that the BLM RecQ helicase has a specific role in early HIV-1 infection.

We assessed effects of SLX4 (FANCP) and RAD51 knockdown in MDMs on HIV-1 integration. Both SLX4 and RAD51 have been independently shown to interact with early HIV-1 infection (48, 49). SLX4 and RAD51 have competing functions with BLM during DNA repair and fork stabilization (50, 51). SLX4 knockdown did not affect viral integration while RAD51 knockdown resulted in a 2-fold decrease, although a double knockdown of BLM and RAD51 did not further compromise the establishment of HIV-1 provirus (**Fig. S5A**). BLM is one of five human RecQ helicases, a highly conserved subgroup of the superfamily 2 helicases (52). It has been suggested that RecQ helicases have nonredundant and nonoverlapping roles in genome maintenance (53, 54) with shared functional properties stemming from distinct activation and recruitment pathways (55). RECQL1 and RECQL5 were hits in our screen as potential dependency factors for HIV-1. Among the family of RecQ helicases, WRN shares the most structural similarity to BLM. RECQL5 and WRN knockdowns in MDMs did not significantly change HIV-1 integration compared to controls, suggesting a unique activity of BLM in HIV-1 infection (**Fig. S5A**).

The host protein SAMHD1 is important for cell cycle progression and functions in DNA damage responses (56, 57). SAMHD1 exhibits dNTPase activity that regulates cellular levels of dNTPs which restrict HIV-1 infection by depleting pools of dNTPs required for reverse transcription (58). To assess whether BLM knockdown and the resulting restriction in HIV-1 integration was due to SAMHD1-mediated activity, cells were first knocked down for either BLM, SAMHD1 or both BLM and SAMHD1 and assayed for changes in expression by western blot and qPCR (**Fig. 3D**, **Fig. S6A**). BLM knockdown did not alter the expression of SAMHD1 or its phosphorylation at Thr592 in MDMs from multiple donors (**Fig. S7**). Consistent with previous results, we observed increased levels of HIV-1 integration in MDMs knocked down for SAMHD1 (59, 60). SAMHD1 knockdowns resulted in an approximately a 4-fold increase in HIV-1 integration in MDMs from 3 donors (**Fig. 3E**). BLM and SAMHD1 double knockdown resulted in an average 2-fold increase of integration in MDMs compared to siRNA controls. The effect of BLM knockdown in SAMHD1 depleted cells was a 50% reduction of integration compared to SAMHD1 only knockdown (**Fig. 3F**). Together these results indicate that BLM knockdown in MDMs inhibits HIV-1 integration, and this inhibition is independent of SAMHD1-mediated HIV restriction.

### BLM knockdown decreases intact proviral integration

Macrophages and resting T cells are less permissive to HIV-1 infection and show a propensity to generating defective proviruses (61, 62). We explored whether BLM contributes to proviral integrity by calculating the ratio of intact and defective proviruses present in HIV-1 infected MDMs with BLM knockdown. Compared to scramble siRNA control, there was a significant decrease in the percent of intact integrated proviruses in cells with diminished BLM, as measured by intact provirus detection assay (IPDA) droplet digital PCR (ddPCR) (63) (**Fig. 4A, 4D**). The decrease in intact proviruses is due to a loss in signals for *env* containing genomes, as indicated by the increased percentage of 3’ defective proviruses (**Fig. 4B**) and no change in the percentage of 5’ defective proviruses (**Fig. 4C**). We also utilized IPDA ddPCR for MDMs treated with ATM, ATR, and PARP-1 inhibitors and observed significant decreases of intact proviral integration in ATM-and ATR-inhibitor treated cells (**Fig. 4E, 4F**).

**Figure 4.**
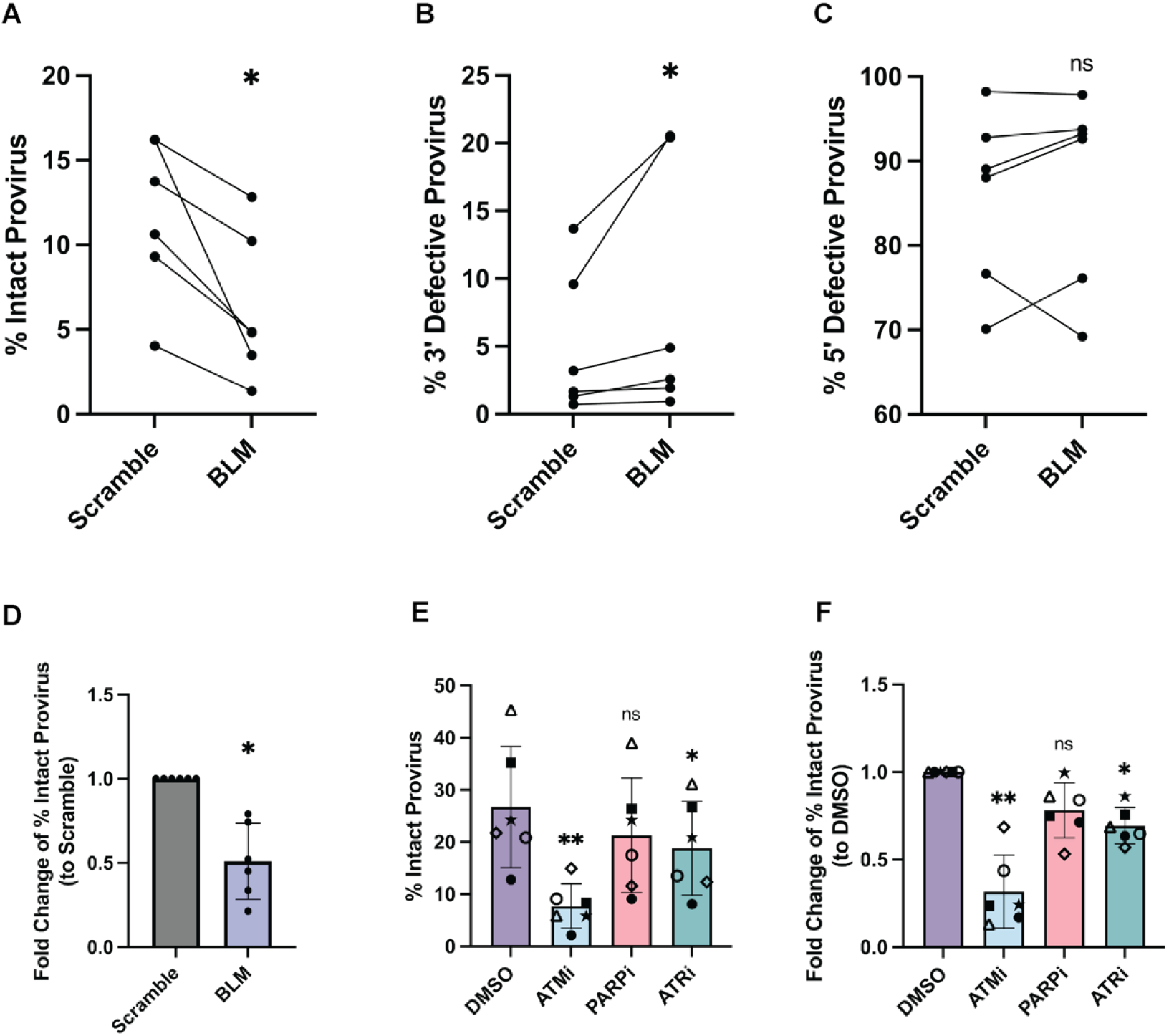
Inhibition of BLM in macrophages decreases intact proviral integration **(A)** Percentage of intact HIV-1 provirus detected in infected MDMs quantified by IPDA. **(B)** Percentage of 3’ defective provirus. **(C)** Percentage of 5’ defective provirus. **(D)** Fold change in percentage of intact HIV-1 provirus (paired) compared to scramble siRNA treated cells. **(E)** Percentage of intact HIV-1 provirus detected in infected MDMs treated with DMSO or DDR pathway inhibitors. **(F)** Fold change in percentage of intact HIV-1 provirus compared to DMSO treated cells. Data are displayed as the means ± SEM and represent ex vivo infection of MDMs from 6 different donors. Statistical significance assessed by Friedman’s test with Dunnet’s multiple comparisons to DMSO control or paired two-tailed t-test (to scramble siRNA control). ns: not significant, *: p<0.05, **: p<0.01

### BLM knockdown inhibits HIV-1 infection at late reverse transcription

The results above indicate a block at or prior to HIV-1 integration in cells with diminished BLM expression. To address if BLM is influencing reverse transcription, we measured levels of RT products in cells with BLM knockdown (**Fig. 5A**). Total viral RT DNA was significantly reduced in BLM knockdowns as assessed by qPCR using primers for late RT products (64), however, there were no significant changes observed for intermediate and early RT products in BLM knockdowns compared to scramble controls. Determining the ratio of these RT products in cells with BLM knockdown showed no change in the ratio from early to intermediate RT products and a significant 2-fold decrease from intermediate to late RT (**Fig. 5B, 5C**). There were no significant differences in RT products in infected cells with WRN and RECQL5 knockdowns (**Fig. S5B-D**). We evaluated viral cDNA levels by measuring 2-LTR circle formation to determine if there were changes in unintegrated episomal HIV-1 genomes following BLM knockdowns (**Fig. 5D**). Using qPCR, a 50% decrease in 2-LTR circles was observed in MDMs with diminished BLM. This decrease was comparable to the decrease in HIV-1 provirus integration and the decrease in total viral cDNA observed following knockdowns. These data are consistent with BLM functioning at late reverse transcription and support a specific non-redundant function for BLM in early HIV-1 infection that is not compensated by other RecQ helicases.

**Figure 5.**
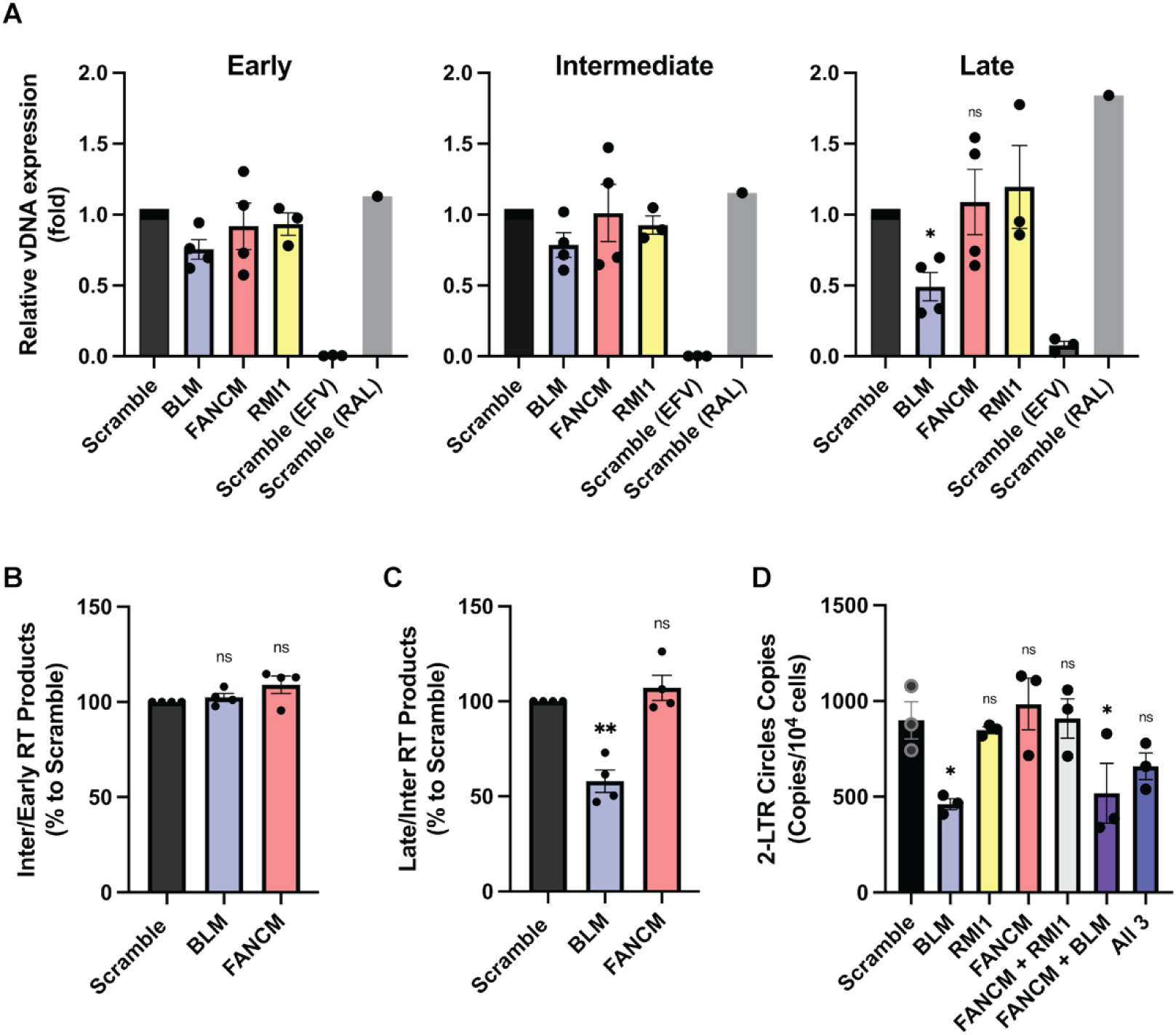
BLM knockdown inhibits HIV-1 infection at late reverse transcription MDMs treated with indicated siRNAs were infected with HIV-1. Controls include cells pre-treated with efavirenz (EFV, 1 μM) or raltegravir (RAL, 30 μM) 1h before infection. **(A)** Total cellular DNA was extracted 2 dpi and used to measure early, intermediate, and late viral reverse transcription products by qPCR. **(B)** Percentage of intermediate RT products and **(C)** late RT products were determined by comparing ratios to scramble siRNA control. **(D)** 2-LTR circles were measured by qPCR. Data are displayed as the means ± SEM with each point representing a different MDM donor. Statistical significance assessed by 1-way ANOVA with Dunnet’s multiple comparisons (to scramble siRNA control). ns: not significant, *: p<0.05, **: p<0.01.

### BLM binds HIV-1 DNA during reverse transcription

Our results indicate BLM is required for efficient reverse transcription in macrophages. The ability and affinity of BLM binding to DNA species with ssDNA/dsDNA junctions led us to hypothesize that BLM binds reverse transcribed HIV-1 DNA by stabilizing the second strand transfer viral DNA structure and facilitating the completion of reverse transcription. To investigate if BLM binds HIV-1 DNA during reverse transcription, we infected CHME3 cells with HIV-1 and performed Cleavage Under Targets and Release Using Nuclease (CUT&RUN) followed by qPCR to detect RT-BLM intermediates. The primers used for this experiment spanned HIV-1 DNA across the U5 LTR, primer binding site (PBS), psi, or the central polypurine tract (cPPT). To ensure HIV DNA-protein binding signal did not result from interactions between targeted proteins and integrated proviral DNA, infections in the presence of integrase inhibitor, raltegravir, were included. We observed a significant enrichment of HIV-1 DNA signal from CUT&RUN-qPCR with anti-BLM and anti-RT pulldowns, when compared to non-specific IgG controls (**Fig. 6A-D**). Increased signal was observed across U5, PBS and psi HIV-1 targets (**Fig 6A-C**). There was no discernable HIV-1 DNA signal from RAD51, H3Kme4 or IgG pulldowns. No HIV-1 DNA signal was detected in pulldowns with uninfected cells or HIV-1 infected cells treated with RT inhibitor efavirenz (**Fig. 6F**). Together these data demonstrate the ability of BLM to bind HIV-1 DNA, before proviral integration and support a model in which BLM interacts with late RT DNA intermediates to facilitate complete reverse transcription and integration.

**Figure 6.**
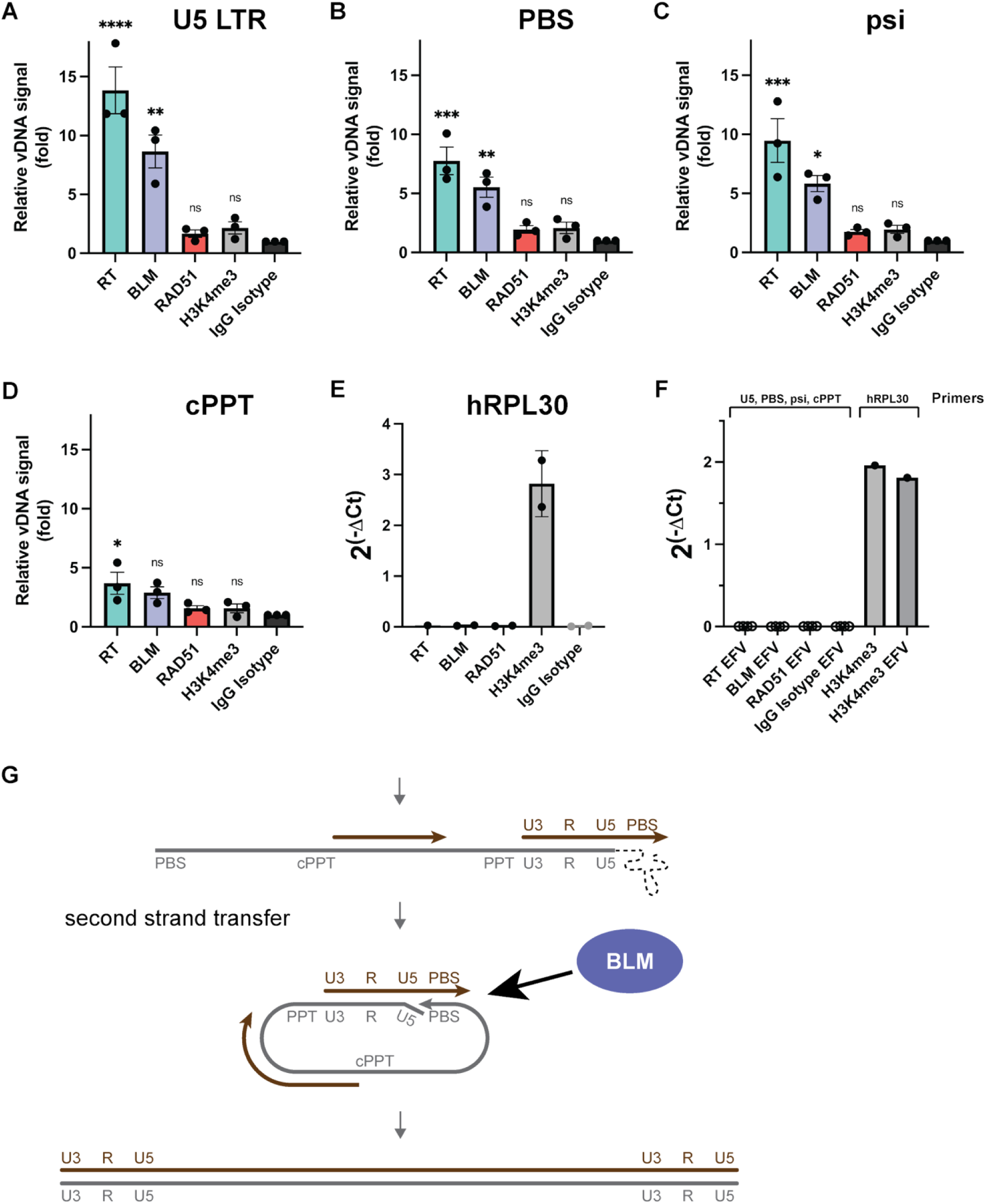
BLM binds HIV-1 DNA during reverse transcription CHME cells were infected for 6 h with NL4-3 BaL (MOI 1) in the presence of raltegravir (RAL, 30 μM) before fixation for CUT&RUN. Protein-bound DNA pulldown with indicated antibodies measured using primer sets corresponding to HIV-1 **(A)** U5 LTR, **(B)** primer binding site (PBS), **(C)** psi, and **(D)** central polypurine tract (cPPT) regions during reverse transcription by qPCR. Samples were normalized to input and IgG isotype negative control. **(E)** Protein-bound DNA from cells with indicated antibody pulldown used to measure control hRPL30 signal by qPCR. **(F)** Infected CHME cells treated with efavirenz (EFV, 1 μM) (except for “H3K4me3”) before fixation. qPCR primers for indicated DNA targets are shown. Data are displayed as the means ± SEM with each dot representing an independent pulldown experiment. Statistical significance assessed by 1-way ANOVA with Dunnet’s multiple comparisons (to IgG control). ns: not significant, *: p<0.05, **: p<0.01, ***: p < 0.001, ****: p < 0.0001. **(G)** Model depicting BLM facilitating completion of reverse transcription at the second strand transfer step by binding the structured single-stranded DNA/double-stranded DNA products.

## Discussion

HIV-1 integration into the host genome necessitates DDR protein interactions. The breadth of host DNA damage responses that assist early infection remain unclear. In this study, we establish that inhibition of ATM and ATR pathways in primary monocyte-derived macrophages results in reduced HIV-1 infection, corroborating previous findings performed in other cell types (22, 38, 65). While both ATM and ATR inhibition restricted HIV-1 integration, only ATM inhibition resulted in a reduction of reverse transcription products and intact proviruses. Using a DDR-specific CRISPR screen we identified DDR factors that influence HIV-1 infection including BLM, which also had a critical role in HIV-1 reverse transcription and the generation of intact HIV-1 proviruses in macrophages, suggesting BLM is downstream of an ATM response initiated during HIV-1 infection.

Our screen identified a total of 137 genes that potentially influence HIV-1 infection and hits included DDR genes previously implicated in HIV-1 infection. For example, our screen identified IDH1, MRE11, PSMA7, RMI1, DDOST, DHX15 and TOP3B which were also hits in a genome-wide siRNA screen by Konig *et al*. (66). Our CRISPR screen also identified several Fanconi anemia repair pathway factors including FANCI, FANCD2, BRCA2 (FANCD1), FANCB, FANCC, FANCG, FANCM, SLX4 (FANCP), RAD51 (FANCR), UBE2T (FANCT) and downstream polymerase REV3L (POLZ). Previous studies have shown Fanconi anemia repair factors, in particular FANCI and FANCD2, facilitate HIV-1 integration (15). Our results highlighting Fanconi anemia factors underscore the importance of this pathway in HIV-1 infection as well as provides confidence in our screening results. Most importantly, we show BLM is a key DDR factor that mediates HIV-1 infection, including RT and integration.

BLM has roles in DNA repair and Holliday junction dissolution that are largely mediated by the BTR complex consisting of BLM helicase, topoisomerase IIIa (TOP3A) and RecQ-mediated genome instability proteins 1 and 2 (RMI1 and RMI2); all except TOP3A were detected in our screen. However, our results suggest that BLM facilitates HIV-1 reverse transcription independently of the other BTR components at the second strand transfer step. Furthermore, other RecQ helicases did not influence HIV-1 integration and reverse transcription indicating a unique function for BLM in HIV infection.

Our screen also identified the recombinase RAD51 as a potential HIV-1 restrictive factor. It has been reported that activating RAD51 restricts HIV-1 integration upon infection (49, 67) while others have shown inhibiting RAD51 restricts HIV-1 infection (68, 69). Previous studies have also suggested BLM displaces RAD51 on DNA both in vitro and in vivo (70, 71). The anti-recombinase activity of BLM appears to function on inactive ADP-bound RAD51 filaments (72, 73). However, we were unable to detect RAD51 on reverse transcribed HIV-1 DNA suggesting that RAD51 and BLM do not have competing activities at the late RT step. In our experiments, RAD51 knockdown in MDMs resulted in reduced HIV-1 integration (**Sup Fig. 5A**). The enigmatic role for RAD51 in HIV-1 infection of macrophages still needs to be determined.

We consistently see a two-fold decrease in all steps downstream of RT, consistent with RT being the primary bottleneck that BLM is acting at. It is noteworthy that in a cell-free in vitro reverse transcription system, reverse transcription is completed at an efficiency of approximately 39% in the absence of other host factors (74). The addition of cell lysates enhances reverse transcription completion to almost 50%, highlighting the ability of the RTC to independently complete reverse transcription and the potential contributions of host factors to facilitate the reverse transcription process.

Given BLM has a high affinity for structured DNA at doubled stranded DNA/single stranded DNA interfaces, and our current understanding of HIV-1 reverse transcription, we envision a model where BLM stabilizes and resolves viral DNA structures formed during the second strand transfer of reverse transcription (**Fig. 6G**). The ability of BLM to bind HIV-1 DNA at this step could serve to bridge other host factors during PIC formation and mediate more efficient reverse transcription and subsequent integration.

## Materials and Methods

Experimental details and methods can be found in *<u>SI Appendix</u>*, including sources of cells and reagents, and protocols for virus production and infection. Details of assays using DNA and RNA and CUT&RUN experiments are described in *SI Appendix*. CRISPR-Cas9 screening protocol and analysis are also described in *SI Appendix*.

## Acknowledgments

We thank Brian Tilton at the BUMC Flow Cytometry Core Facility and assistance provided by Providence/Boston CFAR Scientific Core (P30 AI042853). A.A.L was funded in part by the Immunology Training Program T32 AI007309. In addition, the project was supported by National Institutes of Health grants R01 AI187175, R01 DA055488, R01 DA059952 to AJH & SG.

## SUPPLEMENTAL MATERIALS

### Materials and Methods

#### Cells

CHME3 cells (also known as HMC3: ATCC# CRL-3304)(1), HEK293T cells (ATCC# CRL-3216) and TZM-bl cells (NIH AIDS Reagent Program) were cultured in Dulbecco modified Eagle medium (DMEM; Invitrogen) supplemented with 10% fetal bovine serum (FBS; Gemini Bio-Products) and 1% penicillin/streptomycin (pen/strep; Invitrogen). Peripheral blood mononuclear cells (PBMCs) were isolated from leukapheresis packs (New York Biologics Inc and Rhode Island Blood Center) using Ficoll-Paque Premium (Cytiva, 45-001-751). CD14+ monocytes were isolated from PBMCs by positive selection using the EasySep Human CD14 Positive Selection Kit II (STEMCELL Technologies). Monocyte-derived-macrophages (MDMs) were generated by differentiating CD14+ monocytes for 5-6 days in Roswell Park Memorial Institute medium (RPMI; Invitrogen) supplemented with 10% heat-inactivated Human AB Serum (Millipore Sigma), 1% pen/strep) and 20 ng/mL recombinant human M-CSF (BioLegend) at 37°C and 5% CO_2_.

#### Plasmids

HIV-1 proviral clones HIV-1 NL4-3 and HIV-1 NL4-3 BaL *env* were obtained from the AIDS Reagents Program. The GFP reporter HIV-1 used for DDR-CRISPR screens is a single-cycle HIV-1 encoding GFP in place of Nef (HIV-1/LaiΔenv GFP). HIV-1 packaging plasmids psPAX2 (Addgene, #12260),) and VSV-G envelope expression construct, pMD2.G (Addgene, #12259 were used as previously described (2). Lentiviral CRISPR knockout plasmids were purchased from Addgene as a pool containing DDR-targeting sgRNAs that were cloned into lentiCRISPR v2 backbone (Addgene #52961). DNA Damage Response MKOv4 library was a gift from Dr. Junjie Chen (Addgene #140219). DDR MKOv4 plasmids were amplified to maintain 1000-fold coverage and prepared for lentivirus production as follows: In replicates of 4, 2 μL of 50 ng/μL DDR MKOv4 library was electroporated into 25 μL of ElectroMAX Stbl4 cells (Invitrogen, #11635-018). Cells were recovered in 975 μL SOC medium. 8 mL of recovered cells were pooled and plated on 35 15-cm LB agar plates (in house) containing 100 mg/mL carbenicillin (Gibco, # 10177012), based on dilution plate calculations to maintain 1000-fold coverage of siRNAs (4,530,000 colonies for 4530 sgRNAs). Colonies were pooled, pelleted, and plasmids harvested with plasmid Maxiprep (Invitrogen, #K210017).

### Viruses

#### Virus Generation

Lentiviruses were generated by transfection of HEK293T cells as previously described (3). Briefly, HEK293T cells were co-transfected with 10 μg of either HIV-1 NL4-3, HIV-1 NL4-3 BaL *env*, or HIV-1/LaiΔenv GFP plasmids with 1 μg of VSV-G envelope plasmid, using polyethyleneimine (Sigma-Aldrich, #765090) or Lipofectamine 3000 (Invitrogen, # L3000015) transfection reagent. After approximately 16 hr, cells were washed once with PBS and replenished with fresh media before harvesting supernatants after 24-48 hr and treating with DNase I (50U/mL) (Promega, Z3585) for 1 h at 37°C, in the presence of 10mM MgCl_2_. Supernatants were passed through a 0.45 μm filter (Fisher, # 09-754-21) and lentiviruses were concentrated on a 20% sucrose (Fisher, # S5-500) cushion by ultracentrifugation (24,000 RPM for 1.5 h at 4°C) (SW28 rotor, Beckman Coulter). Virus multiplicity of infection (MOI) was determined with TZM-bl cells as previously described (4).

#### DDR-CRISPR Lentivirus Generation

Lentivirus pool of the DDR MKOv4 library was generated by transfection with Lipofectamine 3000 of HEK293T cells while maintaining 1000x guide coverage as described previously (5,6). Transfected plasmids included 4.8 μg psPAX2, 3.8 μg pMD2.G, and 8 μg DDR MKOv4 plasmid pool.

#### Infections

MDMs were infected with HIV-1 at an MOI of 0.5 or 1, in the presence of 10 μg/mL polybrene (Millipore Sigma, # TR-1003-G) by spinoculation (2,300 rpm, 1.5 h at room temperature), cultured for 1-2 h at 37°C, washed 3x with PBS to remove unbound virus before culturing. CHME cells were infected without polybrene and without spinoculation.

#### Drug treatments

For DDR inhibitors, cells were pretreated with ATM inhibitor KU-55933 (5-10 μM; Millipore Sigma, #SML1109), ATR inhibitor VE-821 (5-10 μM; Millipore Sigma, #SML1415) or PARP inhibitor Olaparib (5-10 μM; Millipore Sigma, #SML3705) for 18-24 h at 37°C prior to infection. DMSO (Sigma, #D8418) was used as a vehicle control. For the indicated experiments, cells were pretreated with efavirenz (1 µM; NIH AIDS Reagent Program) or raltegravir (30 µM; Selleck Chemicals) for 30 min at 37°C prior to infection

### siRNA transfections

#### Primary MDMs

MDMs were transfected with SMARTPool siRNAs (Horizon) (**Supplemental Table 1**) at 50 nM on day 0, then again at 50 nM on day 1 using Lipofectamine 3000 transfection reagent in Opti-MEM. 72 h post-transfection cells were infected as described above.

#### Immortalized Cell Lines

HEK293T cells were transfected with SMARTPool siRNAs (Horizon) (**Supplemental Table 1**) at 40 nM for 48 h using Lipofectamine 3000 transfection in Opti-MEM.

#### RNA isolation and RT-qPCR

Total mRNA was isolated using Quick-RNA Miniprep kit (Zymo Research, #R1054) and reverse-transcribed using oligo(dT)_20_ primer and Superscript III RT (Invitrogen, #18080-044). cDNA corresponding to 100 ng of RNA was analyzed by qPCR using SYBR green (ABclonal, #RK21203) to quantify host mRNA levels and HIV transcripts. C_T_ values of target mRNA were normalized to GAPDH mRNA (ΔC_T_), then ΔC_T_ of the target mRNA was further normalized to control sample by the 2-^ΔΔC^_T_ method.

### DNA isolation and quantitation

Total DNA was isolated from cells by ammonium acetate-isopropanol precipitation. RT products and 2-LTR circles were quantified by qPCR using SYBR green (ABclonal, #RK21203). Primers for early, intermediate, and late RT products and GAPDH controls were used as described (7). Primers for unintegrated viral DNA were used to detect and quantify 2-LTR circles as previously described (8,9). The integrated viral DNA was quantitated by nested Alu-PCR as described previously (10,11). Data were normalized to a standard curve generated from HIV-1 NL4-3 plasmid and genomic DNA from singularly integrated ACH-2 cells (AIDS Reagent Program).

### Western Blot analysis

Cells were lysed in 100 µL of RIPA lysis buffer supplemented with protease and phosphatase inhibitor cocktail (Thermo Scientific, #78442). Protein concentration was assessed via Bradford Assay (Thermo Scientific, #23238) and lysates (20 µg or 40 µg) were combined with 6x loading dye prior to gel electrophoresis. Lysates were loaded in Novex tris-glycine mini protein 4-20% gels (Invitrogen, #XP04200BOX) and run at 120V for 1-2 h. Protein samples were separated with SDS-PAGE on 4-20% polyacrylamide gels (Invitrogen, #XP04205BOX) and transferred onto PVDF membranes (MilliporeSigma, #IPVH00010). Membranes were blocked with Pierce™ Protein-Free Blocking Buffer (Thermo Scientific, #37572) for 1 h at room temperature, followed by overnight incubation with primary antibodies at 4°C. Washes were performed in TBS-T. Secondary antibodies were incubated for 1 h at room temperature and signal was detected with LiCor Odessey scanner.

Blots were performed with the following antibodies: rabbit anti-BLM (Bethyl Laboratories, #A300-110A, 1:1000), rabbit anti-BLM (Abcam, #ab2179, 1:1000), rabbit anti-RMI1 (Proteintech, #14630-1-AP, 1:1000), rabbit anti-FANCM (Proteintech, #12954-1-AP, 1:1000), rabbit anti-RAD51 (Cell Signaling Technology, #8875S, 1:1000), rabbit anti-SAMHD1 (Cell Signaling Technology, #76700S, 1:1000), rabbit anti-phospho-SAMHD1 (Cell Signaling Technology, #89930S, 1:1000), goat anti-rabbit IgG secondary antibody Dylight 680 (Thermo Fisher, SA5-35518, 1:10,000), and goat anti-rabbit IgG secondary antibody Dylight 800 (Thermo Fisher, SA5-35571, 1:10,000). Membranes were scanned using an Odyssey scanner (Li-Cor).

### ELISA

96-well non-sterile ELISA plates were coated with 100 υL/well of HIV-Ig (NIH AIDS Research and Reference Reagent Program, #3957) diluted in PBS (50 mg/mL) and blocked with 200 μL/well of 5% FBS in PBS. Serial diluted p24^gag^ recombinant protein (Advanced BioScience Laboratories, Inc. Lot #B-53) was used as a standard. p24^gag^ was bound with primary antibody p24^gag^ (AIDS Reagent Program, #1513), secondary antibody 100 μL/well of goat anti-mouse-HRP (Sigma, #A2554-1ML). and detected with TMB ELISA substrate (Thermo Scientific, #34021). Absorbance was read at 450 nm.

### Cleavage Under Targets & Release Using Nuclease DNA/Protein binding assay

CUT&RUN experiments were performed following manufacture protocols (Cell Signaling Technology, #86652). CHME cells were infected with HIV-1 NL4-3 BaL at an MOI of 1, for 6 h. Pretreatment of EFV or RAL was done 1 hr before infection. Cells were harvested and fixed in a final concentration of 1% formaldehyde (Cell Signaling Technology, #12606). For each sample, input was collected prior to fixation. Nuclei were bound to concanavalin A beads and permeabilized with digitonin. Samples were incubated with control antibody Tri-Methyl-Histone H3 (Lys4) (C42D8) Rabbit mAb (Cell Signaling Technology, #9751), control antibody rabbit (DA1E) mAb IgG XP® Isotype (Cell Signaling Technology, #66362), rabbit polyclonal anti-HIV-1 RT antibody (Abcam, #ab63911), rabbit polyclonal anti-BLM (Bethyl Laboratories, #A300-110A), or rabbit mAb anti-RAD51 (Cell Signaling Technology, #8875S). Following antibody binding, extracts were incubated with pAG-MNase, 1 mM calcium chloride and then the reaction was stopped and cross-linking reversed using 0.1% final SDS (Cell Signaling Technology, #20533) and 20 μg/mL proteinase K (Cell Signaling Technology, #10012). DNA, including input samples, was purified using ChIP DNA Clean & Concentrator kit (Zymo Research, #D5205). qPCR was performed using primers designed to target HIV-1 U5 LTR, primer binding site (pbs), psi and cPPT regions. Control primers targeted human RPL30 gene (Cell Signaling Technology, #7014) and GAPDH (Thermo). (all primers are in **Supplemental Table 2**)

### DDR-CRISPR pooled screen

CRISPR-KO screens using a targeted DDR-specific sgRNA library were performed by following the Moffat lab protocol Revision 20190723 (6), with modifications for HIV-1 infection.100,000 CHME cells were infected with the pooled sgRNA lentivirus at an MOI of 0.3, for each condition, in triplicate, in the presence of 8 μg/mL polybrene. After 24 h cells were selected on puromycin for 48 h to enrich for cells expressing the DDR sgRNA library and Cas9. Timepoint 0 (T0) aliquots of cells were harvested and frozen at −80°C for sequencing prep. Cells were then infected (except “no infection” controls) with HIV-1/LaiΔenv GFP. 2 dpi cells were sorted into GFP positive and GFP negative populations. Cells were cultured for 5 days before genomic DNA was harvested and prepared as previously described (2) for Illumina NextSeq 500 sequencing. Genomic DNA was isolated by ammonium acetate salt precipitation and quantified by UV spectroscopy. The library was amplified using a two-stage PCR approach. For PCR1 amplification, samples were divided into 50 μL PCR reactions. Each well consisted of 25 μL of NEB Q5 High-Fidelity 2X Master Mix (New England Biolabs, #M0494X), 2.5 μL of 10 μM forward primer Nuc-PCR1_Nextera-Fwd Mix, 2.5 μL of 10 μM reverse primer Nuc-PCR1_Nextera-Rev Mix and 20 μL of genomic DNA (5 ug each reaction). PCR1 cycling settings: initial 30 s denaturation at 98°C; then 10 s at 98°C, 30 s at 65°C, 30 s at 72°C for 25 cycles; followed by 2 min extension at 72°C. PCR1 samples were concentrated by isopropanol precipitation and normalized to 20 ng/μL. Each PCR2 reaction consisted of 25 μL of NEB Q5 High-Fidelity 2X Master Mix (New England Biolabs, #M0494X), 2.5 μL 10 μM Common_PCR2_Fwd primer, and 2.5 μL of 10 μM reverse i7 indexing primer. PCR2 cycling settings: initial 30 s at 98°C; then 10 s at 98°C, 30 s at 65°C, 30 s at 72°C for 13 cycles. PCR products were again purified by SPRI (Beckman Coulter, #B23318), pooled and sequenced on an Illumina NextSeq 500 at the Azenta NGC Laboratory (GENEWIZ) using standard Nextera sequencing primers and 75 cycles. Data were processed by the Data Science Core at Boston University Chobanian & Avedisian School of Medicine. FASTQ files were processed and trimmed to retrieve sgRNA target sequences, followed by comparisons of sgRNAs in the reference sgRNA library file using MAGeCK (12).

## Supplemental Figures

**Fig S1.**
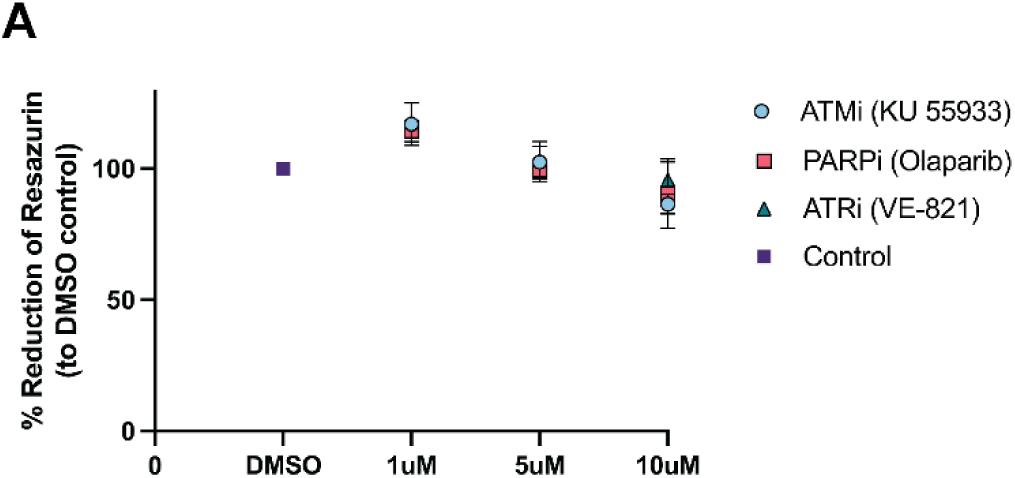
Cell viability with DDR inhibitors **(B)** MDMs were treated with DMSO (10 μM) or KU-55933 (ATM inhibitor), olaparib (PARP1 inhibitor), or VE-821 (ATR inhibitor) at indicated concentrations for 72 h. PrestoBlue cell viability reagent was added to media (1:10) and incubated for 1 h before absorbance was read at 570nm and 600nm. Percentages calculated and compared to normalized DMSO control.

**Fig S2.**
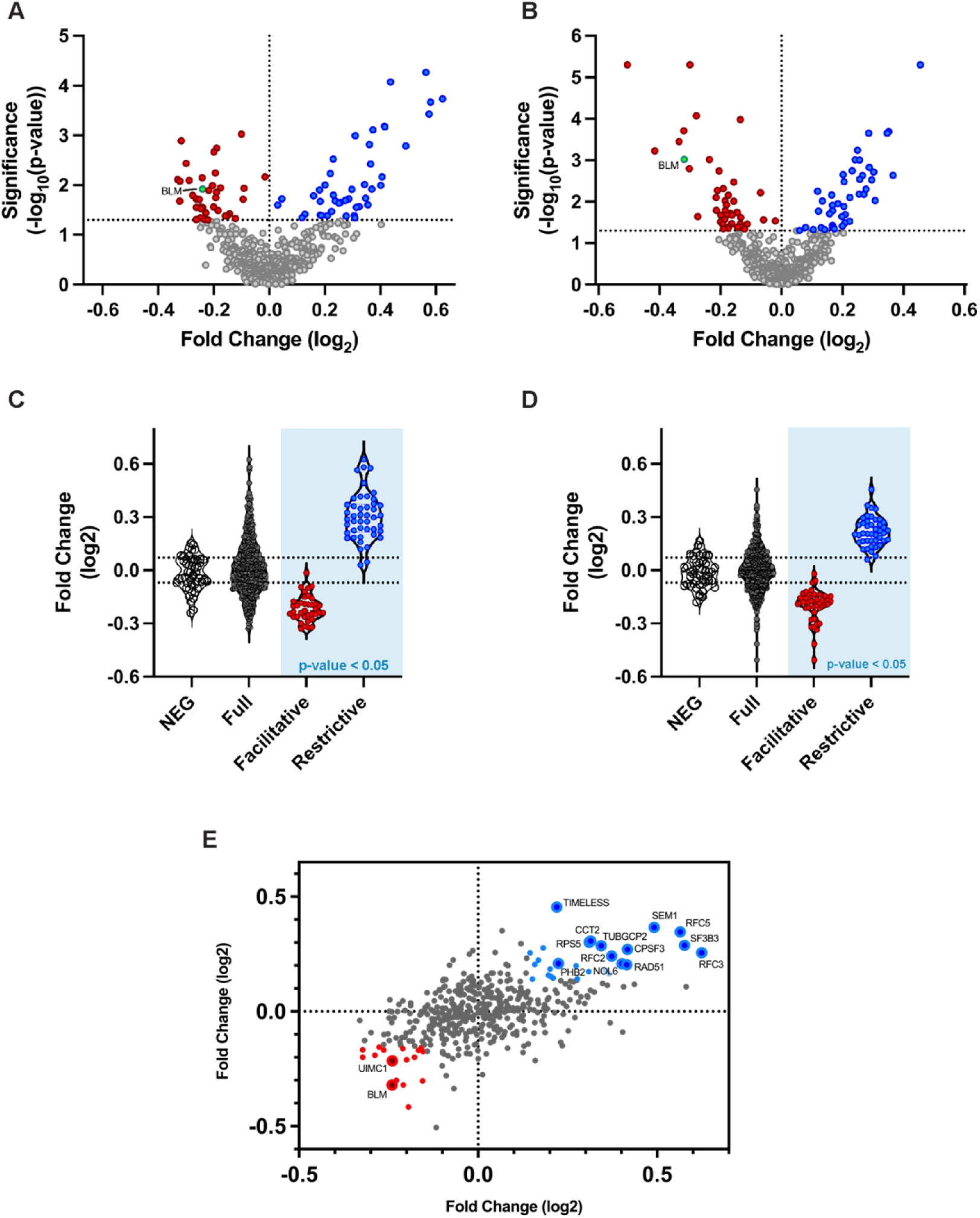
CRISPR Screen identifies DDR genes influencing HIV-1 infection **(A)** Volcano plot for gene hits generated by plotting -log10(p-values) over log2 fold change (LFC) for GFP-positive cells against GFP-negative cells (**supplemental data set 1**) **(B)** Volcano plot for gene hits comparing GFP-positive cells against uninfected cells (**supplemental data set 2**). **(C)** Violin plots of LFC for GFP-positive cells against GFP-negative cells and **(D)** and GFP-positive cells against uninfected cells. Y-axis lines represent ± 5% LFC expression. **(E)** Scatterplot comparing log2 fold change of both analyses (vs GFP-negative and uninfected cells). Labeled genes represent enriched (blue) or depleted (red) genes in both analyses greater than 15% LFC.

**Fig S3.**
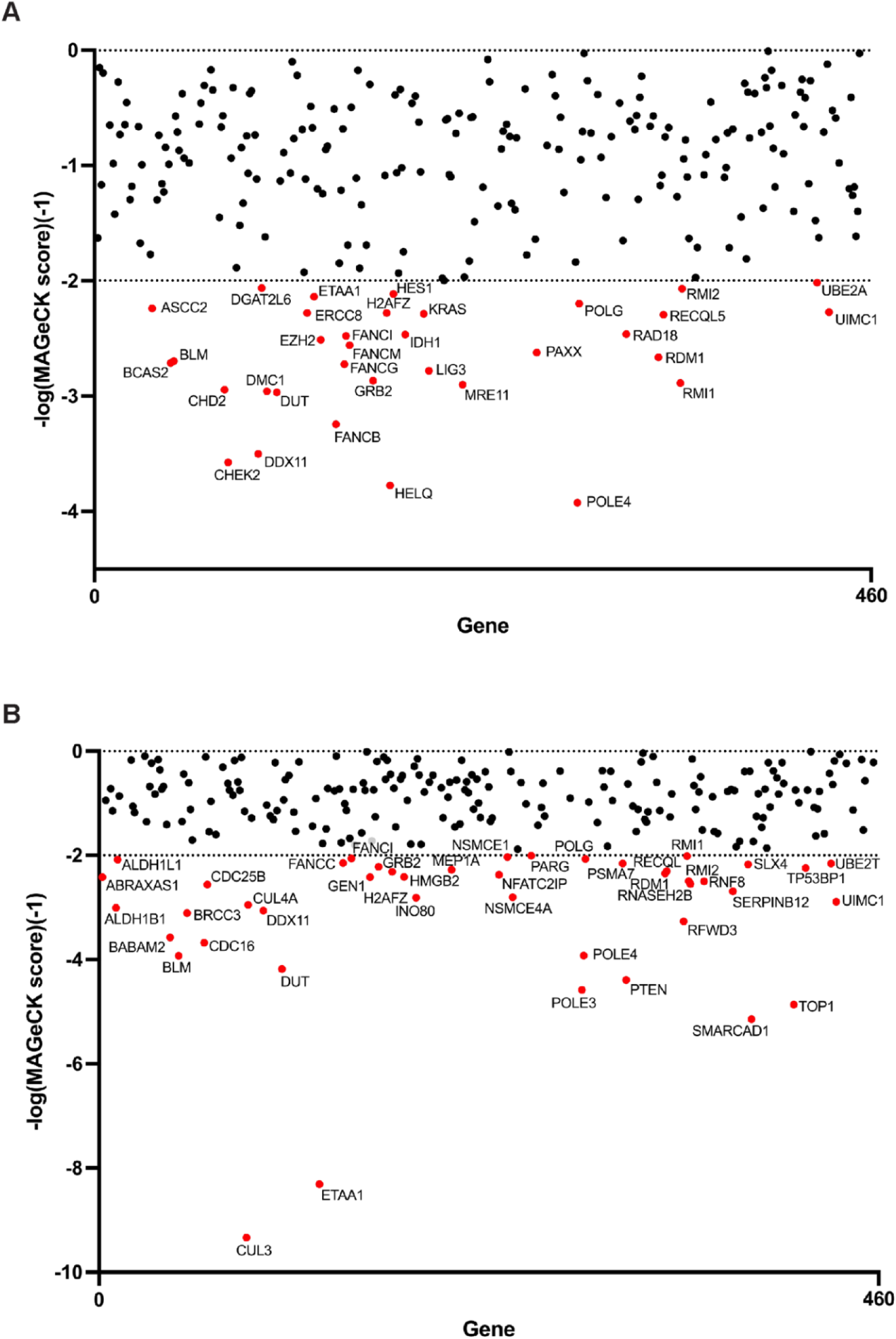
Genes identified that potentially facilitate HIV-1 infection Scatterplot of all genes depleted in analyses comparing GFP-positive cells against **(A)** GFP-negative cells and against **(B)** uninfected cells.

**Fig S4.**
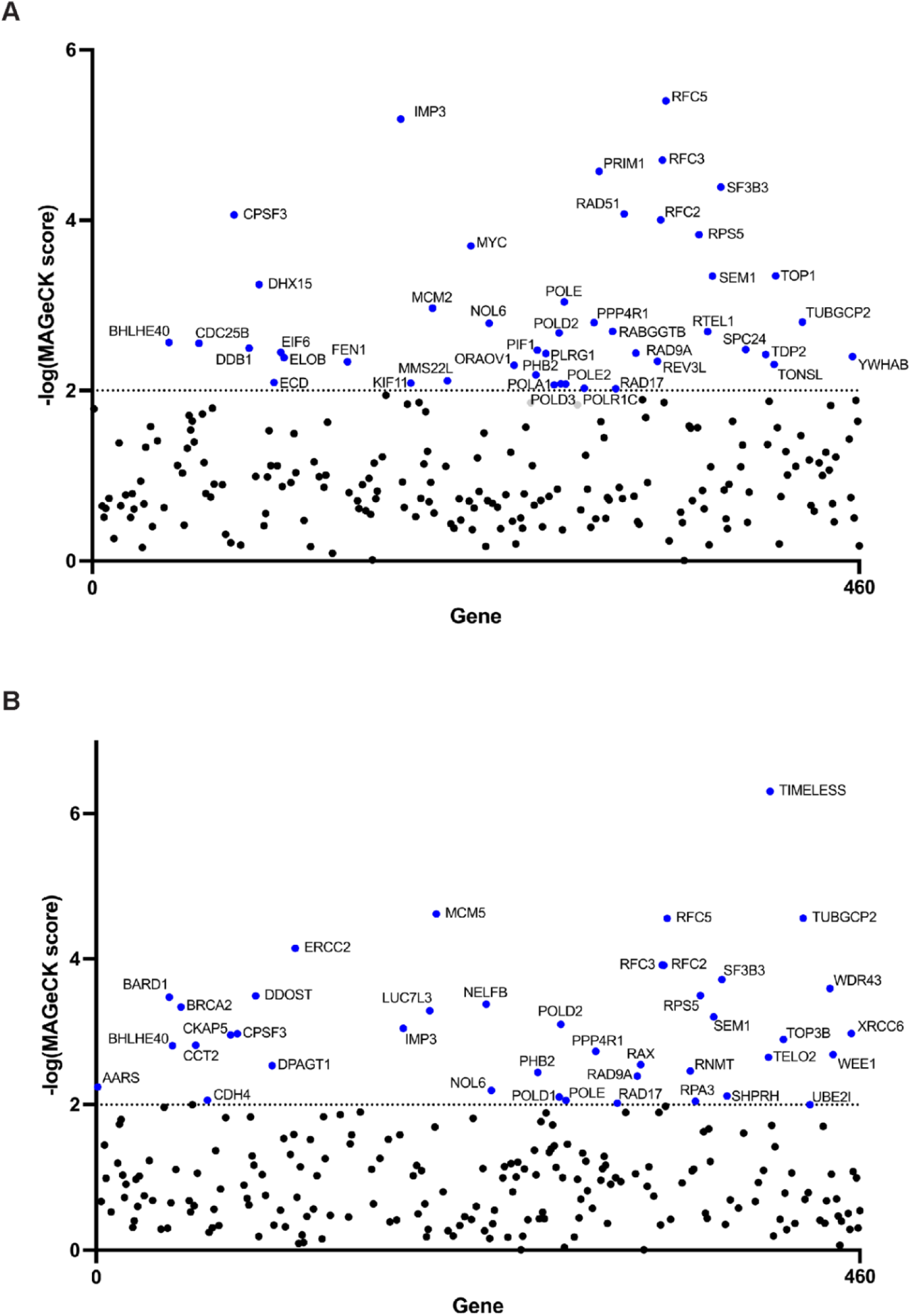
Genes identified that potentially restrict HIV-1 infection Scatterplot of all genes enriched in analyses comparing GFP-positive cells against **(A)** GFP-negative cells and against **(B)** uninfected cells.

**Fig S5.**
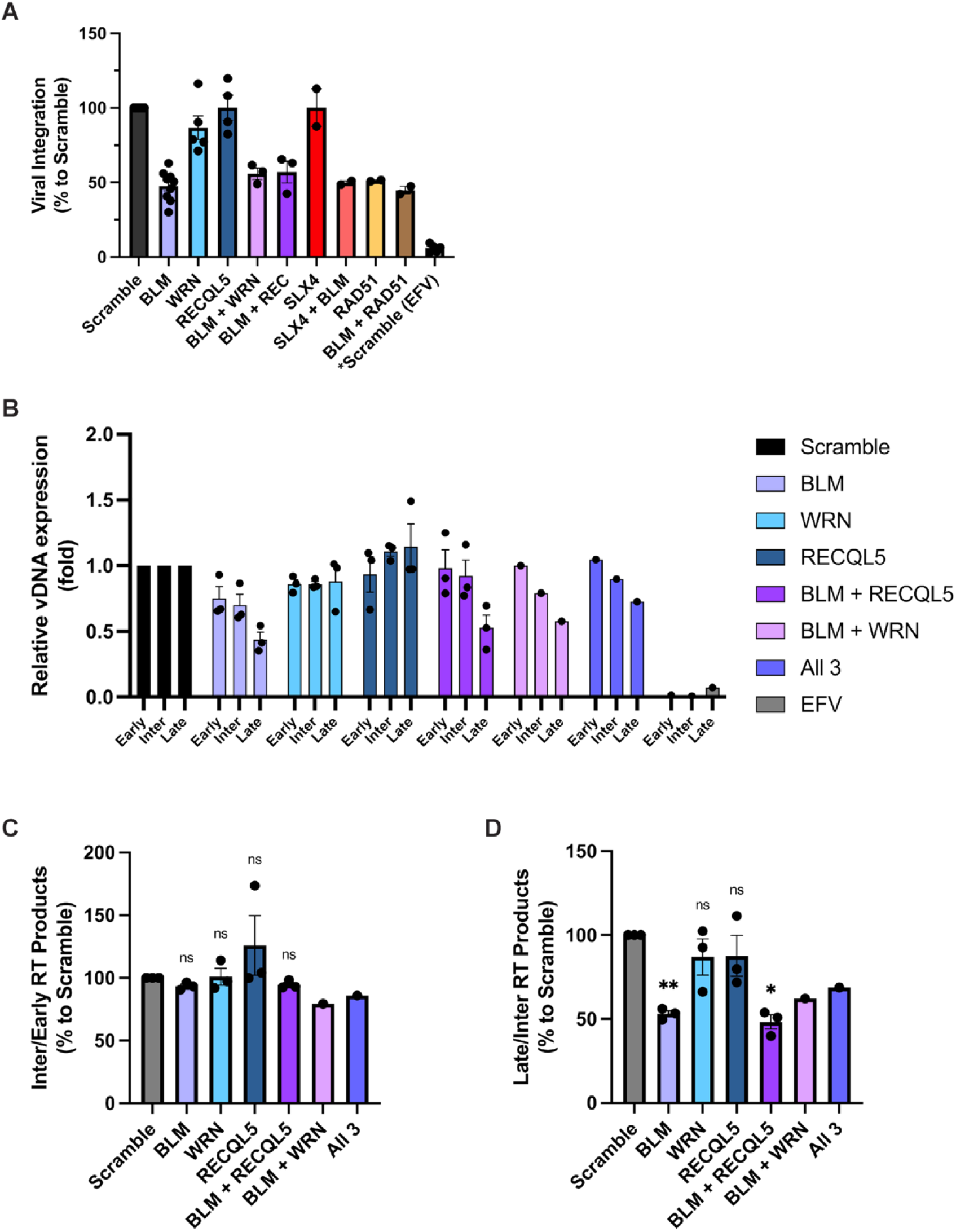
Assessing knockdown of DDR factors in MDMs during HIV-1 infection **(A)** MDMs were infected with NL4-3 BaL after pre-treatment with siRNAs and 2 dpi, proviral integration was measured by Alu-PCR. **(B)** Early, intermediate, and late viral reverse transcription products were measured by qPCR. **(C)** Percentage of intermediate RT products and **(D)** late RT products were determined by comparing ratios to scramble siRNA control. Data are displayed as the means ± SEM with each point representing a different MDM donor. Statistical significance assessed by 1-way ANOVA with Dunnet’s multiple comparisons (to scramble siRNA control). ns: not significant, *: p<0.05, **: p<0.01.

**Fig S6.**
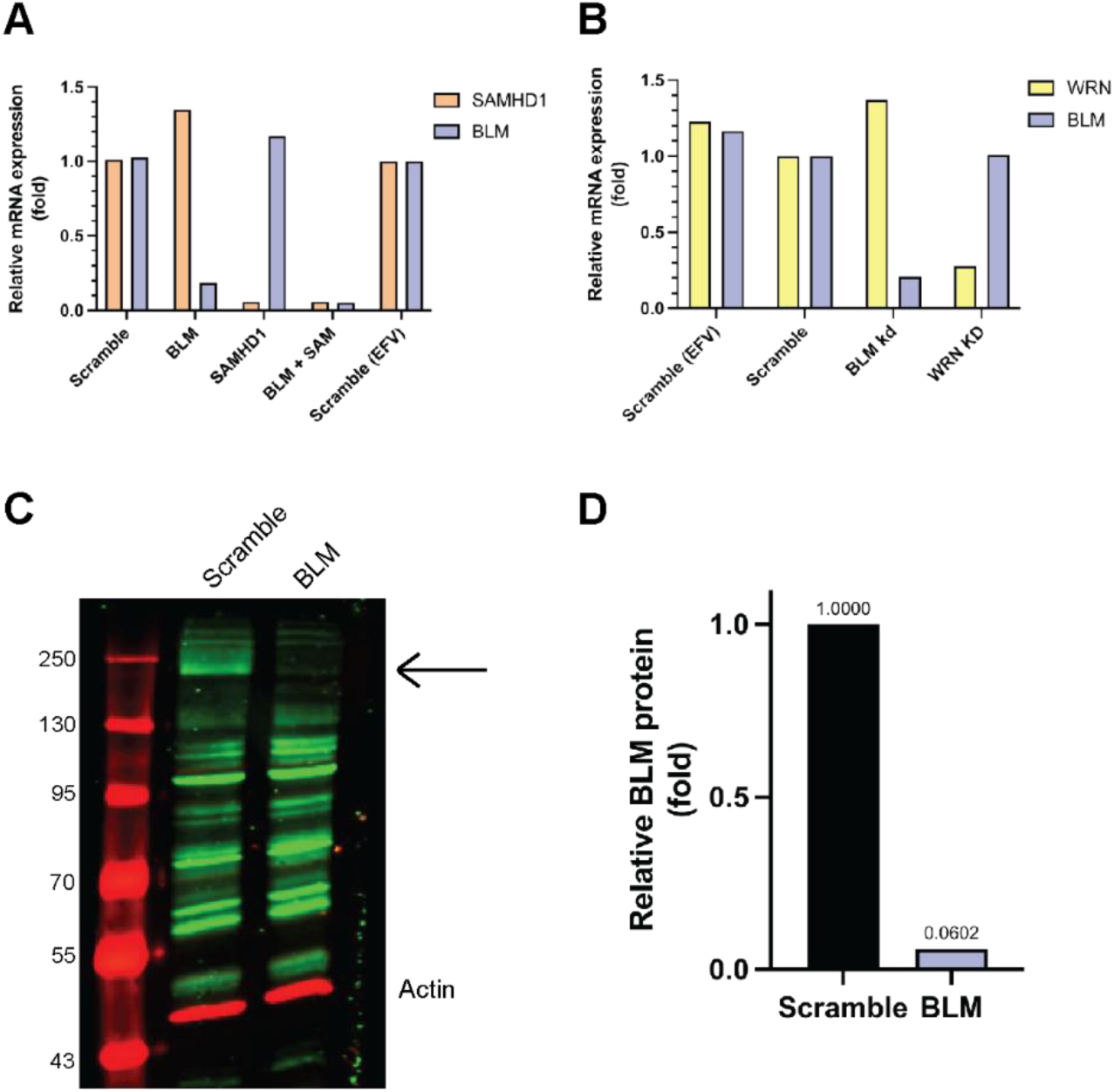
mRNA knockdown correlates with protein knockdown. **(A)** mRNA expression for each targeted gene indicated was measured by qPCR following a double-treatment of 100 nM siRNAs in MDMs. **(B)** mRNA expression for each targeted gene indicated was measured by qPCR following treatment of 40 nM siRNAs in HEK293T cells. **(C)** siRNA knockdown of BLM in HEK293T cells was examined by western blot **(D)** and quantified. Figure represents n = 1 data averaged from 1-2 million HEK293T cells per siRNA knockdown run in technical triplicate for qPCR.

**Fig S7.**
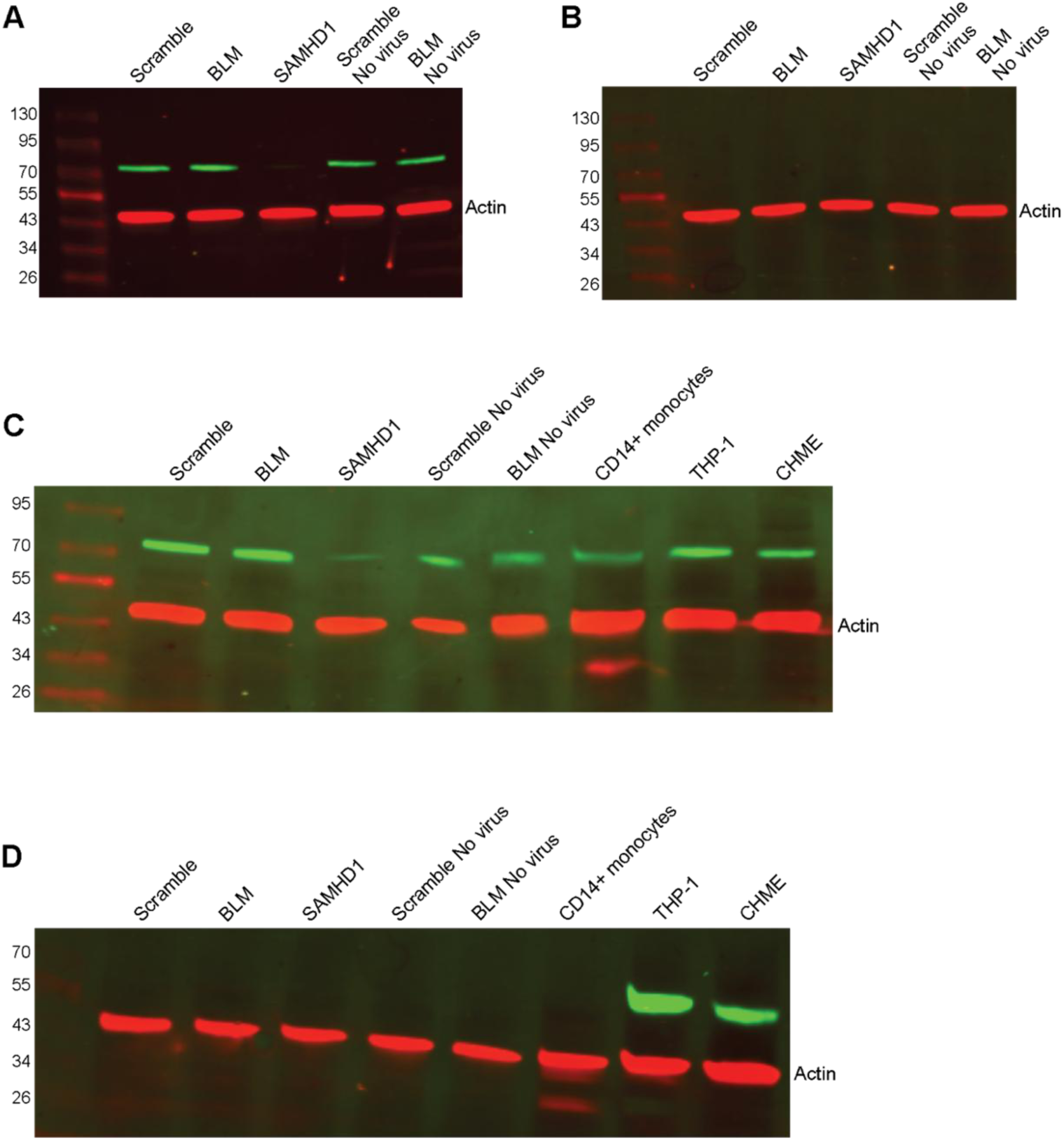
BLM knockdown does not effect SAMHD1 expression and SAMHD1 phosphorylation **(A)** Expression of SAMHD1 (green) assessed by Western blot in MDMs infected with NL4-3 BaL after pre-treatment with siRNAs. **(B)** Expression of phosphorylated SAMHD1 at Thr592 (green) assessed by Western blot. **(C)** Samples expanded to include uninfected primary CD14+ monocytes, THP-1 cells, and CHME cells for SAMHD1 expression and **(D)** phosphorylated SAMHD1 at Thr592.

---

**Table S1.**
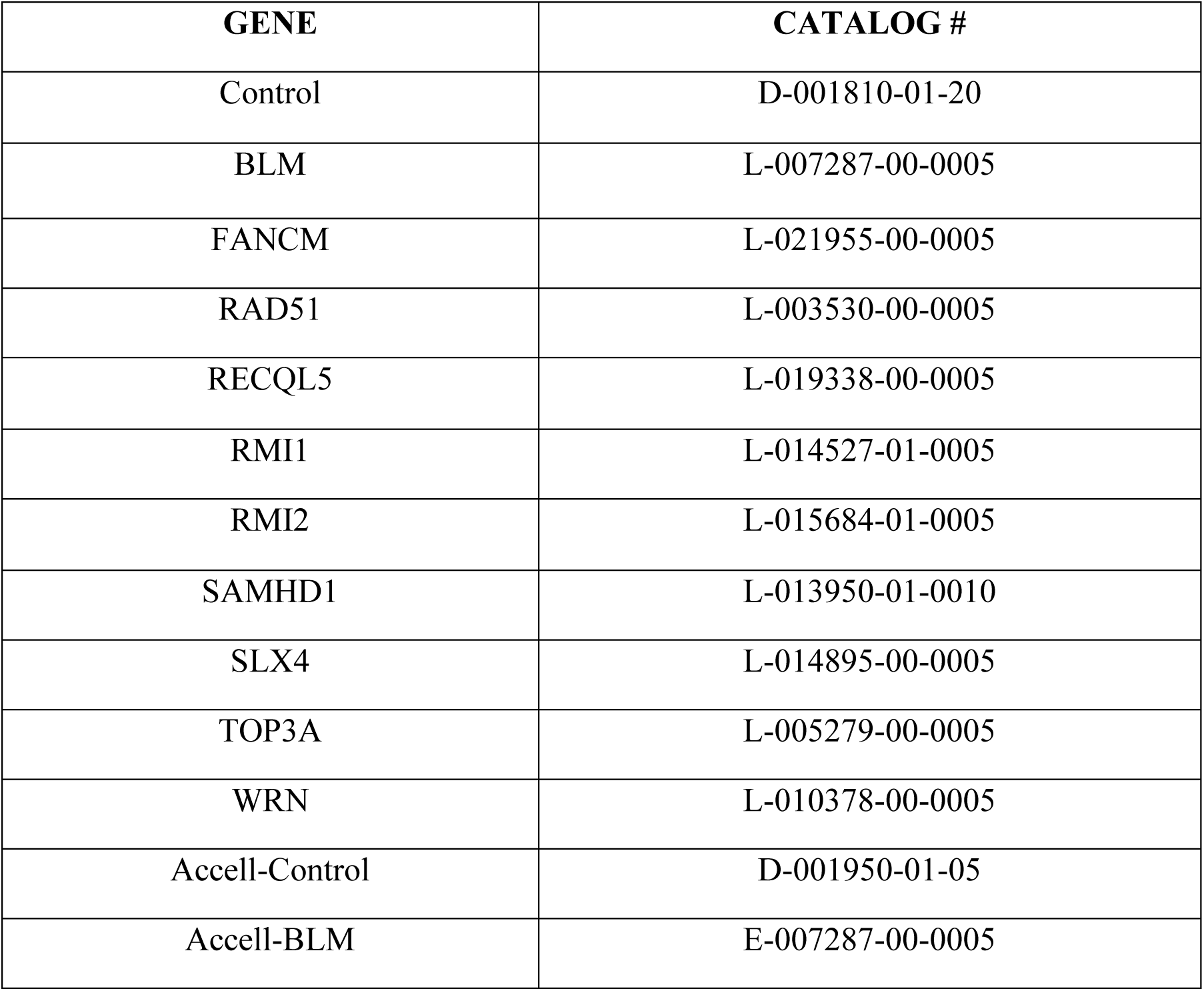
SmartPool siRNA Catalog Numbers.

**Table S2.**
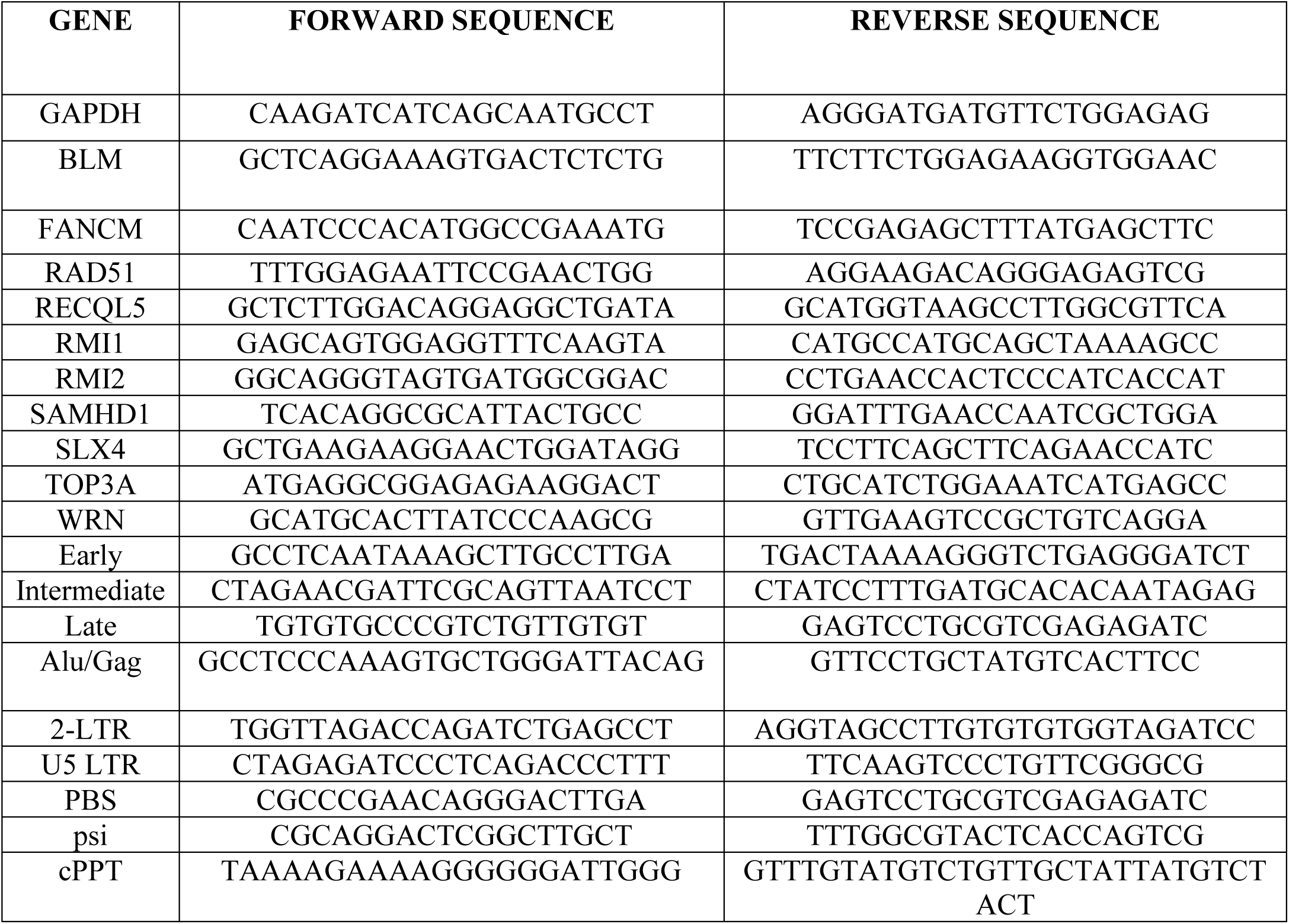
qPCR Primer Sequences.

